# Rapid expansion of synaptic complexity as a key contributor to cognitive growth in early humans

**DOI:** 10.1101/2025.04.09.648028

**Authors:** Guilherme Lepski, Analía Arévalo, Patricia Silva de Camargo, Kelly Nunes, Renan Barbosa Lemes, Tiago Ferraz, Andre Strauss, Shigeru Miyagawa

## Abstract

Primates have been evolving for over 50 million years. At some point, humans made an unusually large evolutionary leap, giving rise to abilities like the creation of tools, intricate art and complex language. The neuronal synapse, a key player in information processing and brain plasticity, has largely been ignored as a potential factor in this process. Here we used the genomic databases of ancestral hominins to compare the expression levels of 995 genes expressed in the human nervous system among archaic (6 Neanderthal, 2 Denisovan) and modern humans (62 African Modern Human). We searched in the 95^th^ top p-value for variants whose derived alleles had a frequency ≥90% in modern and <10% in archaic humans. We then used the STRING database to perform protein-protein interaction networks on the 95^th^ top p-value for the variants. We identified genetic variants in 15 genes, and in two (STX16 and UBASH3B), the allele frequency was significantly higher in modern versus archaic humans. These genes have previously been associated with critical cellular (proliferation, differentiation, migration, survival) and synaptic (exocytosis, synaptic vesicle fusion) processes, supporting the idea that changes in synaptic structure and function may have played a key role in the development of human cognition.

## Introduction

Cultural evolution in the human lineage is a matter of great interest among experts of different fields, including archaeology, anthropology, linguistics and neuroscience. Indirect evidence from each of these fields may be combined to tell a story that can begin to explain the relatively rapid leap in human cognitive capacity that gave rise to advanced abilities such as the creation of sophisticated tools, intricate art and complex language. How and when this revolution took place is a matter of great debate [1-13].

How can we explain this cognitive leap? The Integration Hypothesis (IH; [14, 15]) offers one explanation for the evolution of language. It begins with the assumption that in nature, there are two fundamentally distinct systems underlying animal communication. One is seen in the alarm calls of monkeys and apes, such as that of vervet monkeys, which have calls for leopard, eagle, and snake [16]. These are distinct signals each with “meaning” in the sense that the utterances are connected to some occurrence in the real world, such as the appearance of a leopard. The alarm calls are similar, though not identical, to words, and in IH, they are called the Lexical (L) System. In contrast, birdsongs are finite-state patterns without any meaning on their own [17]. They are expressive, in that a bird sings to express a desire, most commonly the desire to mate. In IH, birdsongs are called the Expressive (E) System. The assumption is that in nature, all communication systems are based either on the L or the E system. Furthermore, the L and E systems have existed for millions of years as we see in primates for L and songbirds for E. IH describes humans as having both the L and the E systems. At some point, possibly around 130 kya, these two systems integrated to form language. This gives the appearance that language emerged rapidly around that time, and in a sense it did, but the components that make up its parts have existed for millions of years. Thus, IH addresses the paradox of evolution and language by supposing that the components that make up language have existed for millions of years, but these components integrated to form language only around 130 kya. There are animals that exhibit both systems, such as the gibbons [18, 19], which possess alarm calls like the vervets, and also sing like birds. However, there is no evidence that the two systems in gibbons have integrated. They remain distinct, unlike in humans.

Many factors have been implicated in the development of intelligence, including brain weight or size, number of cortical neurons, encephalization quotient, neuronal packing density, axonal conduction velocity, and interneuronal distance [20]. However, much less attention has been given to the neuronal synapse, the main structure responsible for transmitting information to neighboring cells, and ultimately responsible for the flow of information. The synapse is the center for all kinds of modulation and plastic changes that are the hallmark of human brain complexity.

In this study we revisited the genomic database of ancestral hominins (Allen Ancient DNA Resource, AADR v.54.1) [21]), with special attention to a list of 995 genes expressed in the human nervous system [22], and measured expression level differences among these genes between archaic (6 Neanderthal, 2 Denisovan) and modern humans (62 African Modern Human). To infer a possible selection signal in modern humans, we searched in the 95th top p-value for variants whose derived alleles had a frequency equal to or greater than 90% in modern humans and less than 10% in archaic humans. Also, the protein-protein interaction networks were performed on the 95th top p-value for variants using the STRING database. Our goal was to search for possible genetic variants that would differ between archaic and modern humans and investigate whether any of those would explain aspects of cognitive processing that may underlie some of the differences observed in evolution.

## Material and Methods

### Neuronal gene selection

We selected approximately 1,000 genes from a high-resolution human gene expression analysis database done by in-situ hybridization (ISH) platform and previously made available online (http://human.brain-map.org/ish). These genes are known to be expressed in the human visual cortex (Brodmann’s areas 17 and 18) and midtemporal cortex (Brodmann’s areas 20, 21 and 22) of modern humans (27 men and 19 women) without any known neuropsychiatric illness. These areas were chosen because they are most widely studied and well conserved within species [22].

### Dataset

For the present study, we analysed six Neanderthal, two Denisovan and 62 African Modern Human samples from the curated Allen Ancient DNA Resource (AADR v.54.1) dataset [21]) genotyped for 1.24k single nucleotide polymorphisms (SNPs). Since the introgression of Neanderthal and Denisovan DNA into modern non-African humans is well reported [23, 24], in our analysis we used only samples from modern sub-Saharan African humans (Table S1). The uneven coverage among these genomes is acknowledged and presents itself as a limitation of the study.

### Methods

To assess differences in genetic variants associated with genes expressed in the brain of modern and archaic humans (Neanderthal+Denisovan), we compared the allele frequencies of genes coding neuronal proteins in modern humans (Table S2; [22]). Thus, from the previously selected set of 995 neuronal genes, the R package biomaRt [25] was used to identify the genomic regions of interest and ±50kbp adjacent to it. Of these, 978 genes and/or adjacent regions have SNPs genotyped by the 1.24k SNP array. For this set of SNPs, ancestral and derived alleles were annotated with Ensembl Variant Effect Predictor (VEP) [26]. Thus, using the R package biomaRt [25]), we identified the genic region of interest and ±50kbp adjacent to it. Allelic frequency was inferred only for SNPs with at least 10 genotyped chromosomes.

Then we applied Fisher’s exact test to compare the allele frequency observed between modern and archaic humans. Analyses were performed using the plink v1.9 software [27]). To identify the most relevant genetic variants, we looked at those with p-values in the top 95th percentile of the upper quartile of the distribution. To check whether the observed allele frequency differences for candidate genes between modern and archaic humans also occur in modern populations outside Africa, we searched for their allele frequencies in dbSNP (https://www.ncbi.nlm.nih.gov/snp/).

To infer a possible selection signal in modern humans, we searched in the 95th top p-value for variants whose derived alleles had a frequency equal to or greater than 90% in modern humans and less than 10% in archaic humans.

Finally, the protein-protein interaction networks were performed on the 95th top p-value for variants using the STRING database [28]). Although this database has known limitations, its known strengths are its comprehensiveness and ease of use, as it covers > 10,000 organisms, draws from a wide diversity of data sources, and offers many intuitive interface features including personalization and enrichment detection [28, 29]. Furthermore, the STRING database can integrate and connect protein-protein functional linkage data to previous knowledge, thus allowing users to access organism-wide protein networks [28].

## Results

The Fisher’s exact test in the region of 978 genes encoding neuronal proteins in modern humans identified 15 candidate genes with genetic variants with distinct allele frequencies between modern and archaic humans (Fig. 1). Of these genes, 13 have alleles with a frequency greater than 90% in modern humans and less than 10% in archaic humans (Table 1), with the *UBASH3B* (rs11218809 T>C; p-value=3.04^-16^) and *STX16* (rs4812024p-value=1.20^-10^) being the derived alleles with a higher frequency in modern humans. More details regarding the candidate alleles, their functionality and frequency in other modern human populations can be found in Table S3.

**Figure 1.**
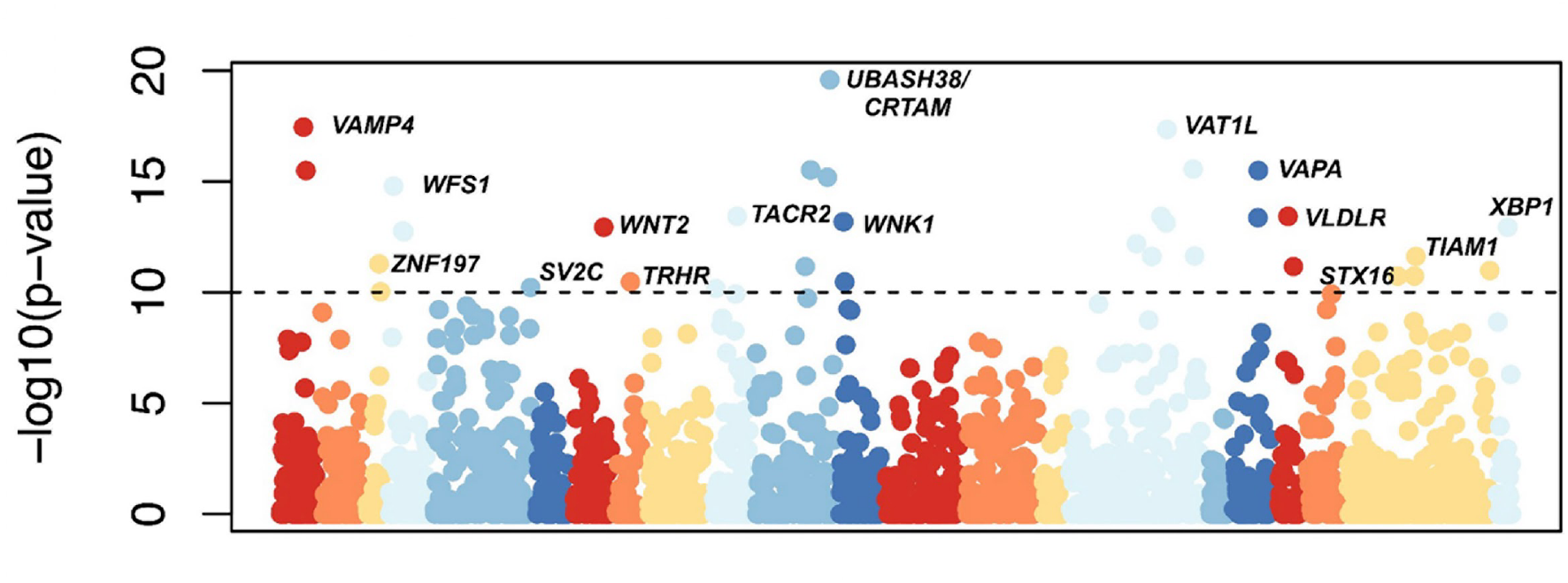
Manhattan plot of Fisher’s exact test. The x-axis corresponds to chromosomes and the y-axis to p-values on a -log10 scale. The dashed line indicates the upper 95th quartile of the distribution. Each dot represents a SNP. Candidate gene names are shown.

**Table 1.**
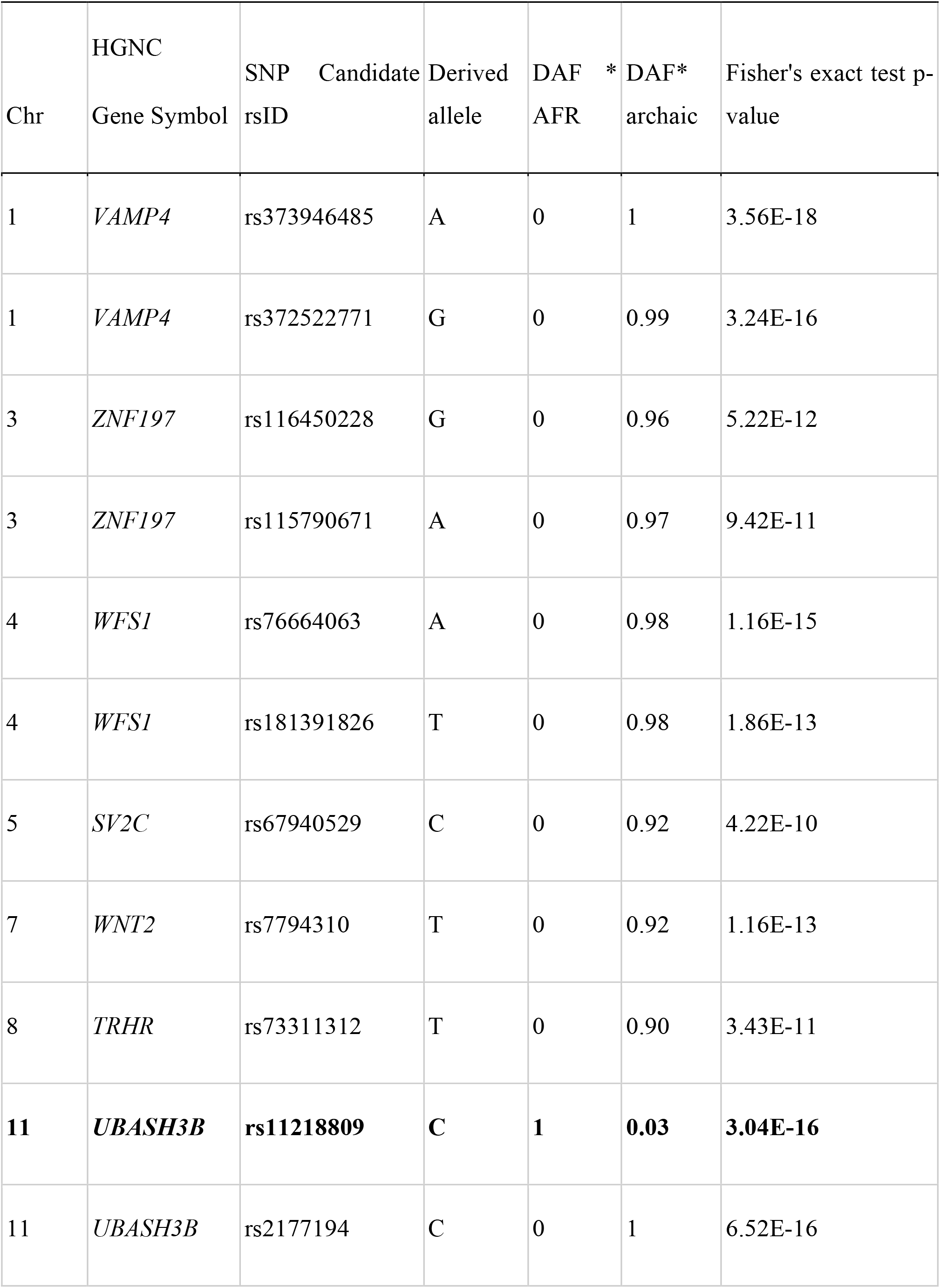
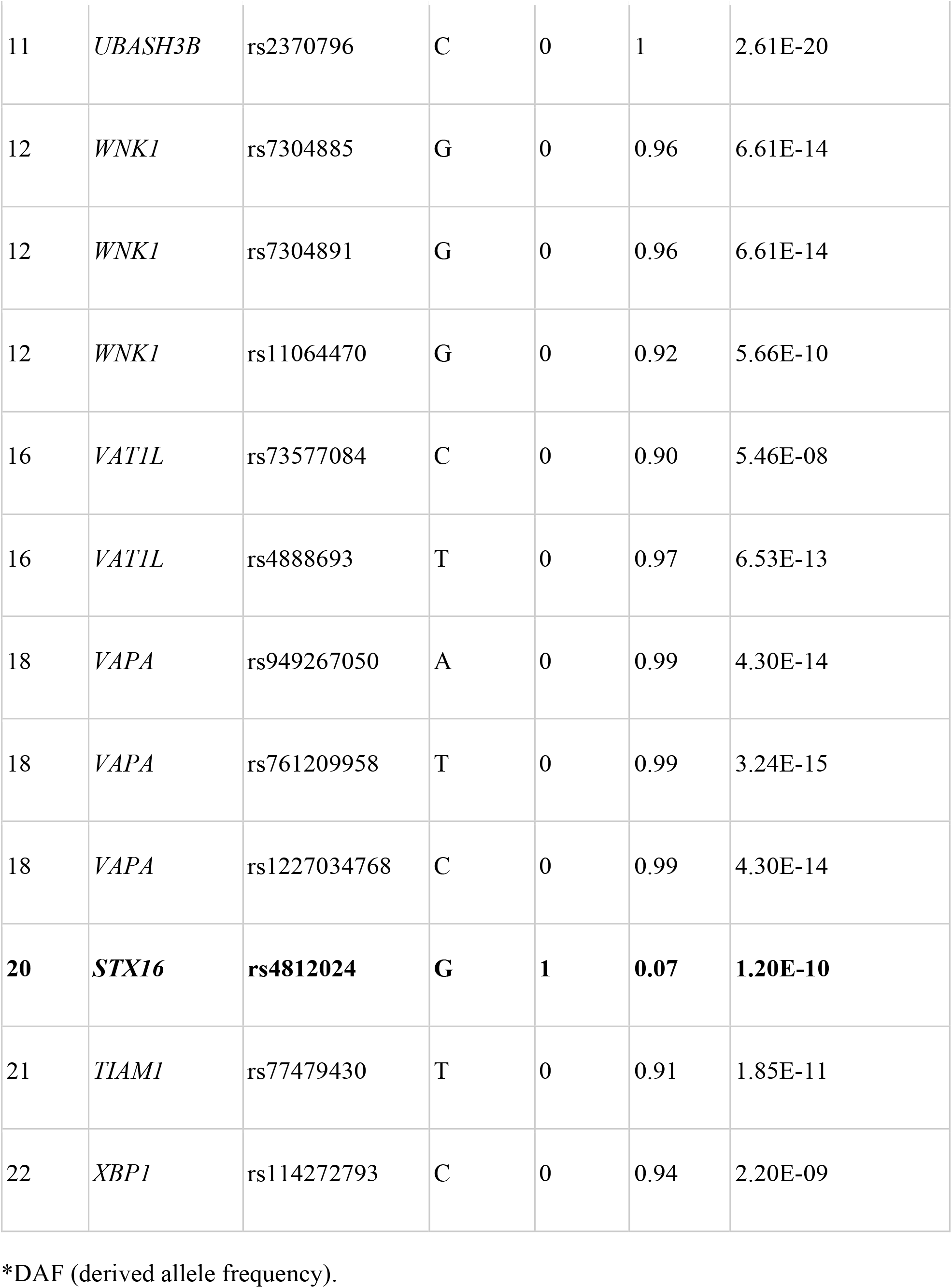
Candidate genes and alleles identified using Fisher’s test. For each of these, we show the derived allele and its frequency in modern African humans and archaic humans.

Next, we produced the protein-protein interaction network among the 13 candidate genes (Fig. 2) and identified two interaction networks: one among the *VAPA, VAMP4* and *STX16* genes and the other between *WSF1* and *XBP1*.

**Figure 2.**
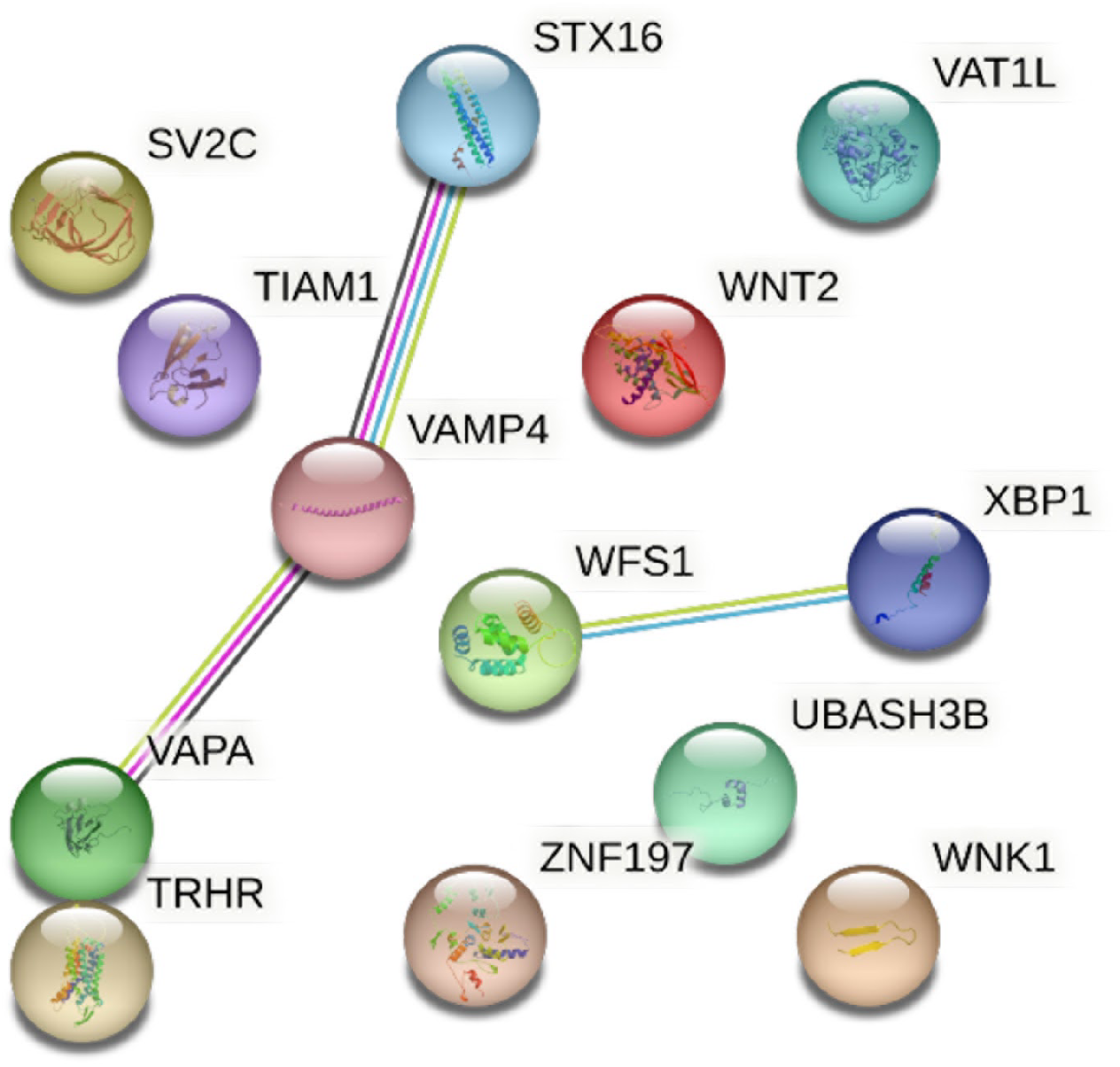
Protein-protein interaction networks. Each sphere corresponds to one of the candidate genes. Line colors indicate the type of interaction evidence: black for co-expression, pink for experimentally-determined, blue for cured dataset and yellow for text mining.

## Discussion

Darwinian or neo-Darwinian theories cannot adequately explain the unexpectedly marked leaps in human intelligence in evolution. Interesting models tried to bring together cultural and genetic evolution [30]. Nevertheless, transmission of cultural traits does not respect the three Mendelian laws of inheritance: segregation, independent assortment, and dominance [31]. From our perspective, this evolutionary jump was only possible because more complex neural networks arose, allowing for higher intelligence levels. If measuring intelligence continues to be one of modern neuroscience’s great challenges, finding supporting evidence within the fields of archaeology and comparative biology must carry even greater difficulties. Previous assumptions, now considered obsolete, were based on brain size or weight, the encephalization quotient [32], or the absolute number of neurons [33]. More recent methods consider multiple variables like number of cortical neurons, neuron packing density, interneuronal distance and axonal conduction velocity—factors that ultimately are supposed to determine general information processing capacity (IPC) [20].

By comparing ∼1,000 genes that code for neuronal proteins in archaic and modern humans, we identified genetic variants in 15 genes. In two of these, STX16 and UBASH3B, the allele frequency was significantly higher in modern than in archaic humans. The UBASH3B gene (ubiquitin associated and SH3 domain containing B) codes an intracellular protein that is highly expressed throughout the entire central nervous system (CNS), but especially so in the cerebellum, cerebral cortex, pons and medulla oblongata. Previous research has reported that the encoded protein inhibits endocytosis of transmembrane receptors, including epidermal growth factor receptor (EGFR) through RPK (receptor tyrosine kinases), thus controlling essential cellular processes like cell proliferation, differentiation, migration and survival [34, 35]. Its universal expression in the CNS suggests a role in delaying the effect of certain neurotransmitters in the synaptic cleft by inhibiting endocytosis. The other relevant gene we observed to be more prominent in modern human species was STX16 (Syntaxin 16). This gene belongs to a family of membrane integrated Q-SNARE (synaptic) proteins that participate in exocytosis. Furthermore, it interacts with VAMP4 (vesicle associated membrane protein) to drive spontaneous and calcium-independent fusion of synaptic vesicles [36].

These data support the idea that evolutionary changes in synaptic structure may have been a key factor in the development of human intelligence. The synapse is a central structure in the brain that can undergo ultrastructural and functional changes induced by activity [37, 38]. This feature (also known as plasticity) is the physiological substrate for learning and memory, and to a certain extent, is also present in all forms of multimodal associations, language, and other higher cognitive functions. In fact, it is the core of the brain’s complexity, as well as the main site of modulation in the nervous system.

How the synapse emerged in evolution is not fully understood [39]. Ancestors of chemical synapses, the so-called Ursynapses, appeared in common ancestors of cnidarians and bilaterians [40]. Historic work on *Caenorhabidities elegans* led to the identification of key synaptic proteins like the one codified by the gene unc-13, which is essential for synaptic vesicle exocytosis [41]. Later studies in vertebrates using proteomics revealed several orthologues with remarkable functional conservation [42]. A more recent study with sponges suggests that the first directed communication in animals may have evolved to regulate feeding, serving as a starting point on the long path toward nervous system evolution [43]. Nevertheless, some diversity arose after two rounds of whole-genome duplication that affected the chordate lineage 150 million years ago [44, 45].

Comparing the synaptic machineries across species is a valuable tool for understanding neuronal complexity in vertebrates [35]. In an interesting study comparing mouse and zebrafish synapse proteomes, Bayés et al. described a higher number of SNARE complex components (Syntaxin, VAMPs and SNAPs) and associated proteins (Synaptotagmins and Sec1/Munc18s) in mice relative to zebrafish brains [46]. These proteins play essential roles in endocytosis, AMPA receptor trafficking, and regulation of synaptic plasticity. By comparing mouse and human DNA by *in situ hybridization* (ISH), Zeng et al. reported 79% conservation between species, but 21% of species-differential expression [22]. Among the significant differences observed were some genes involved in the transcription of ion channels (*SCN4B, KCNC3*), neurofilament (*NEFH*), adhesion molecules (*MFGE8*), but also synaptic proteins (*SYT2 and VAMP1*). This evidence suggests that, despite a certain degree of conservation, synaptic molecular complexity increased consistently, with added functionality and improved fitness [47]. Furthermore, in a recent study targeting the middle temporal gyrus (MTG), an area highly associated with language processing in human, Jorstad and colleagues [48] used deep transcriptomic profiling to compare the cerebral cortex of humans to that of four nonhuman primate species (chimpanzees, gorillas, rhesus macaques and marmosets). They observed that many differentially expressed genes (DEGs) for all primate species were associated with molecular pathways related to synaptic connectivity and signaling. When they then analysed human-accelerated mutations or deletions, they discovered that 15-40% of human-specific DEGs were near genomic regions under adaptive selection [48].

Current understanding of human intelligence, or IPC (information processing capacity) is based on the “continuity theory”, which predicts that humans’ higher cognitive abilities result from conserved evolutionary trends, leading to an increased absolute and relative brain size, as well as more neurons, especially in the frontal lobe [49, 50]. Another line of thinking named “cerebrotype” brain evolution assumes that specific rather than general changes in the brain, particularly in the prefrontal cortex, were responsible for higher cognitive capacity [51, 52]. Important controversies regarding the first theory have recently emerged [20]. A linear model of the relationship between brain volume (also valid for weight) and body size revealed that larger animals have relatively smaller brains, whereas smaller animals have relatively bigger brains. In blue whales, for instance, the brain represents 0.01% of body volume, while in shrews, it accounts for 10% or more of their bodies. In terms of encephalization quotients, a higher EQ in dolphins relative to gorillas is paradoxical, as the latter are considered to be more intelligent than the former. If we consider absolute brain size as the preponderant factor, we should assume that elephants (brain weight 5Kg) or sperm whales (10Kg) are more intelligent than humans (1.35Kg brain, 2% of the body size), and horses (600g) are more intelligent than chimpanzees (273g). One important brain feature is the thickness of the cortical mantle. Whereas in primates it is relatively thick (3 to 5mm), in cetaceans and elephants it is relatively thin (1 to 1.8mm). This causes neuronal packing density (NPD) to reduce in larger animals, and absolute neuronal counts in the cortex to be relatively constant across these mammals (15 billion in humans, as opposed to 10-12 billion in cetaceans and elephants). But the most relevant paradox in these neuronal features arises when we compare human brains to those of certain birds (parrots and corvids). Despite having high NPDs, these birds’ brains are very small. Noteworthy, they do not even possess a cerebral cortex. In birds, the nidopallidum, a part of the telencephalon, is the centre of executive functions, and hodologically equivalent to the primate neocortex, albeit much less complex.

The research we have presented suggests that the concept of intelligence and its neuronal defining features must be revisited. We identified genetic variants in 15 genes, and in two – STX16 and UBASH3B – the allele frequency was significantly higher in modern versus archaic humans. These genes have previously been associated with critical cellular and synaptic processes, including exocytosis, fusion of synaptic vesicles, as well as cell proliferation, differentiation, migration and survival. These findings support the idea that evolutionary changes in synaptic structure may have represented a key aspect in the development of human intelligence, thus driving attention to a structure that until now has been mostly neglected and shifting the search for the origins of intelligence in a new, exciting direction. Additional studies are necessary to support this idea and thus renew the understanding of our own past.

## Acknowledgements

The research of Guilherme Lepski, Analía Arévalo, Patricia Silva de Camargo, and Shigeru Miyagawa was funded by the São Paulo Research Foundation (FAPESP) (grant no. 2018/18900-1), research project “Innovations in Human and Non-Human Animal Communities.”

## Data Availability

The data used for the analysis of this paper are available as supplementary material.

## Author Contributions

**Guilherme Lepski:** Conceptualization, Methodology, Writing - Original Draft. **Analía Arévalo:** Conceptualization, Methodology, Writing - Original Draft. **Kelly Nunes:** Methodology, Formal analysis. **Renan Barbosa Lemes:** Methodology, Formal analysis. **Tiago Ferraz:** Methodology, Formal analysis. **Andre Strauss:** Visualization, Methodology. **Patricia Silva de Camargo:** Visualization, Writing - Review & editing. **Shigeru Miyagawa:** Methodology, Writing - Original Draft, Supervision.

